# Gene Expression: edgeRun: an R package for sensitive, functionally relevant differential expression discovery using an unconditional exact test

**DOI:** 10.1101/008409

**Authors:** Emmanuel Dimont, Jiantao Shi, Rory Kirchner, Winston Hide

## Abstract

**Summary:** Next-generation sequencing platforms for measuring digital expression such as RNA-Seq are displacing traditional microarray-based methods in biological experiments. The detection of differentially expressed genes between groups of biological conditions has led to the development of numerous bioinformatics tools, but so far few, exploit the expanded dynamic range afforded by the new technologies. We present edgeRun, an R package that implements an unconditional exact test that is a more powerful version of the exact test in edgeR. This increase in power is especially pronounced for experiments with as few as 2 replicates per condition, for genes with low total expression and with large biological coefficient of variation. In comparison with a panel of other tools, edgeRun consistently captures functionally similar differentially expressed genes.

**Availability:** The package is freely available under the MIT license from CRAN (http://cran.r-project.org/web/packages/edgeRun)

## 1 INTRODUCTION

Next generation sequencing technologies are steadily replacing microarray-based methods, for instance transcriptome capture with RNA-Seq (Mortazavi et al, 2008) and CAGE-Seq capture for the promoterome (Kanamori-Katayama et al, 2011). All of these approaches result in digital expression data, where reads or tags are sequenced, mapped to the genome and then counted. The discrete nature of the data has required the development of new bioinformatics tools for their analysis that address discrete count data.

Once the expression has been quantified, an important next step is the statistical significance testing of differential expression between two or more groups of conditions. By the far the simplest and most popular approach reduces differential expression to a pairwise comparison of mean parameters, resulting in a fold-change measure of change and a p-value to ascertain statistical significance of the finding. To address this problem, tools such as *edgeR* (Robinson et al, 2010), *DESeq2* (Love et al, 2014) among many others have been developed and can be applied to any experiment in which digital count data is produced.

This vast array of tool choices can be bewildering for the biologist since it is generally not clear under which conditions a tool is more appropriate than its alternates. Traditional metrics used when benchmarking methods such as the false positive rate and power are useful but limited as they are purely statistical concepts that can only be tested on simulated data. Moreover they do not help in determining to what extent methods deliver truly biologically important genes. This is a major challenge because in the vast majority of cases, we do not know what the true positives and negatives are.

In this paper, we propose a novel metric to determine the number of functionally relevant genes reported by a differential expression tool and present *edgeRun*, an extension of the *edgeR* package delivering increased power to detect true positive differences between conditions without sacrificing on the false positive rate. We show using simulations and a real data example that *edgeRun* is uniformly more powerful than a host of differential expression tools for small sample sizes. We also demonstrate how even though it may be less statistically powerful than *DESeq2* in some simulation cases, *edgeRun* nonetheless produces results that are functionally more relevant.

## 2 METHODS

### 2.1 edgeRun: exact unconditional testing

Assuming independent samples, Robinson et al. (2011) proposed *edgeR*, an R package that eliminates the nuisance mean expression parameter by conditioning on a sufficient statistic for the mean, a strategy first popularized by Fisher (1925) for the binomial distribution. This leads to a calculation of the exact p-value that does not involve the mean. The advantage of this approach is its analytic simplicity and fast computation, however a key disadvantage is that this conditioning approach loses power, especially for genes whose counts are small.

We propose an alternative more powerful approach which eliminates the nuisance mean parameter via maximizing the exact p-value over all possible values for the mean without conditioning which we call “unconditional *edgeR*” or *edgeRun*. This technique was initially proposed by Barnard (1945) for the binomial distribution. The main disadvantage of this method is the higher computational burden required for the maximization step. On the other hand, the gain in power can be significant. A thorough derivation and comparison of both methods can be found in the Supplementary Methods.

### 2.2 Benchmarking against other methods

The *compcodeR* Bioconductor package (Soneson, 2014) was used to benchmark the performance of *edgeRun* against a panel of available other tools using a combination of simulated and real datasets. *edgeRun* had the highest area under the curve (AUC) of all methods and it maintained a comparable false discovery rate similar to other tools. In terms of power, only *DESeq2* was found to outperform *edgeRun.* For this reason in the next section, we perform a functional comparison only with *DESeq2*. The full results are summarized in Supplementary Methods.

### 2.3 Comparing functional relevance

We propose to compare the genes called significant by various differential expression tools. Figure 1 compares the results of *edgeRun* and *DESeq2* applied to a prostate cancer dataset (Li et al., 2008) using an FDR < 5% cutoff. Out of the 4226 genes reported as differentially expressed, 80% were common to both tools. The highest 500 up- or down-regulated of these consensus genes by fold-change are used as a seed signature. It is reasonable to hypothesize that true differentially expressed genes uniquely reported by a differential expression tool are functionally connected to genes in the consensus group. We use GRAIL (Raychaudhuri et al, 2009) coupled with a global coexpression network COXPRESdb (Obayashi et al, 2013) to assess the relatedness between a gene and the consensus group. As expected, nearly half of these seed genes are correlated with other members of the seed group, meaning that these consensus genes form a tightly connected network. Figure 1 shows that *edgeRun* reports 6.6 times more unique DE genes, and a larger proportion of which are coexpressed with the consensus. This means that the genes reported by *edgeRun* are more likely to be functionally relevant as they are more correlated with the consensus network.

**Figure 1:**
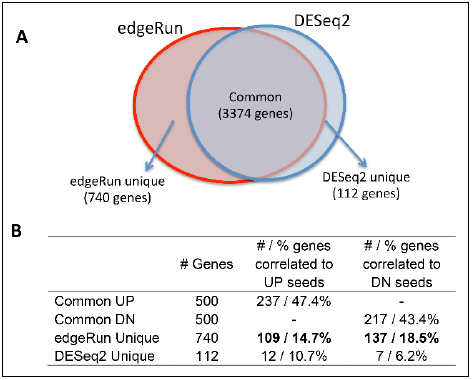
Comparing the functional relevance of genes called significantly differentially expressed by edgeRun and DESeq2

## 3 DISCUSSION

We present *edgeRun*, an R package that improves on the popular package *edgeR* for differential digital expression by providing the capability to perform unconditional testing, resulting in more power to detect true differences in expression between two biological conditions. Even though the computational burden is increased, the power gained using this approach is significant, allowing researchers to detect more true positives, especially for cases with as few as 2 replicates per condition and for genes with low expression, all the while without sacrificing on type-I error rate control. *edgeRun* is simple to use, especially for users already experienced with *edgeR* as it is designed to interface with *edgeR* objects directly, taking inputs and generating output in the same format.

## ACKNOWLEDGEMENTS

We would like to thank Oliver Hofmann, Shannan Ho Sui, Gabriel Altschuler and Yered Pita Juarez for their valuable feedback.

Funding: none.

*Conflict of Interest*: none declared.

